# From Predicted Ki to Surrogate IC50: Similarity-Guided Empirical Calibration of Drug–Target Affinity Predictions

**DOI:** 10.64898/2026.09.04.749433

**Authors:** Basel Mansour, Shayoni dutta, Binil Benny, Sifat Ahmed, Dan Takahashi

## Abstract

Drug–target affinity models return the endpoint on which they are trained, whereas medicinal chemistry decisions are often made with a different assay readout. Here, we trained a DeepPurpose model to estimate inhibition constants (Ki) from molecular graphs and protein sequences and asked whether those predictions could be aligned empirically with measured IC50 values without treating Ki and IC50 as interchangeable. A BindingDB-trained checkpoint retained useful cross-target ranking on the Davis kinase benchmark without Davis training data (concordance index 0.860). We then rebuilt the Ki training set from ChEMBL 37 records coded as Binding assays (assay_type = ‘B’), after removing censored records and targets with poor replicate reproducibility. This reduced median fold error from 16.35× to 9.23× on a leakage-cleaned kinase panel and from 15.30× to 7.90× on a 14-target non-kinase panel before any IC50 calibration. Similarity-guided leave-one-out calibration further reduced the non-kinase panel median error to 3.12× for the original checkpoint and 3.22× for the ChEMBL checkpoint at Tanimoto T = 0.6. Because retraining removed a substantial part of the apparent correction, we interpret the calibration as a target- and chemistry-dependent empirical offset between model output and IC50 assay space, not as a mechanistic Ki-to-IC50 conversion. In a separate project-level stratification of public records, median replicate variability was 2.41× for the subset classified as biochemical Ki and 3.30× for biochemical IC50; a small-sample correction placed the Ki variability nearer 2.8×. These values provide an empirical scale for the remaining calibration error rather than a theoretical performance limit. Similarity, rather than the number of calibrators, governed the main accuracy–coverage trade-off. The resulting values are surrogate IC50 estimates for cross-target triage; within-target ranking remains a limitation of the present architecture.

## 1. Introduction

Early drug discovery is partly a ranking problem. A project can generate or source far more compounds than it can synthesize and test, so computational models are useful only if they help decide which compound–target pairs deserve attention first. Drug–target affinity (DTA) models address this problem by learning a continuous measure of interaction strength from chemical and protein information rather than treating interaction as a simple yes/no event.^1,2^

The practical difficulty is that binding affinity is not a single experimental quantity. Public bioactivity databases contain Ki, Kd, and IC50 measurements collected under diverse assay conditions.^3,4^ Ki is an inhibition constant and, under a defined kinetic model, has a mechanistic interpretation. IC50 is the inhibitor concentration associated with 50% inhibition in a specified assay.^5,6^ For competitive inhibition, Ki and IC50 can be related through the Cheng–Prusoff relationship, which depends on substrate concentration and Km.^5^ The public records assembled for this project did not consistently provide the assay parameters needed to apply that mechanistic relationship record by record. A predicted Ki therefore cannot be relabeled as IC50 without introducing an assumption that the underlying experiment does not justify.

This endpoint mismatch matters because IC50 is a common summary of inhibitor potency in concentration– response analyses.^6^ A DTA model can be useful at the affinity-prediction stage and still be inefficient to connect to downstream potency analysis. We therefore treated Ki prediction and IC50 estimation as two separate problems. DeepPurpose was used to predict Ki from a molecular graph and protein sequence.^1^ A second, empirical layer then estimated the offset between predicted pKi and measured pIC50 from structurally related compounds assayed against the same target.

The study developed iteratively. The first checkpoint was trained on a BindingDB Ki collection. BindingDB is a public resource of experimentally measured protein–small-molecule affinities.^3^ During validation, however, 25.1% of the original training labels were found at exactly pKi 5.00, and the original extract mixed assay contexts. We therefore rebuilt the training data from ChEMBL 37,^4,7^ restricting records to exact Ki values from assays coded by ChEMBL as Binding (assay_type = ‘B’).^8^ We then removed suspect and duplicated entries, filtered targets with poor replicate reproducibility, and collapsed replicate measurements by their median. This second checkpoint provides a useful test of whether calibration is correcting a true endpoint difference or merely compensating for avoidable bias in the underlying affinity model.

Our aim was not to propose another DTA architecture. Instead, we asked whether a standard DTA model could be turned into a more practical potency-estimation workflow by adding a transparent, similarity-aware calibration step. The work was evaluated against an independent kinase benchmark, a leakage-cleaned kinase panel, two dense single-target kinase sets, and fourteen non-kinase targets spanning six pharmacological families. Published Davis results from DeepDTA and GraphDTA are included only as context, because those models were trained directly on the Davis data whereas our tested checkpoint was not.^2,9,10^

## 2. Materials and Methods

### 2.1 Study design

The workflow had two stages (**Figure 1**). Stage 1 predicts pKi from a compound–protein pair. Stage 2 calibrates that value toward pIC50 using measured compounds from the same target. The calibration is evaluated separately from the DTA model so that improvements in endpoint agreement are not mistaken for improvements in ranking.

**Figure 1.**
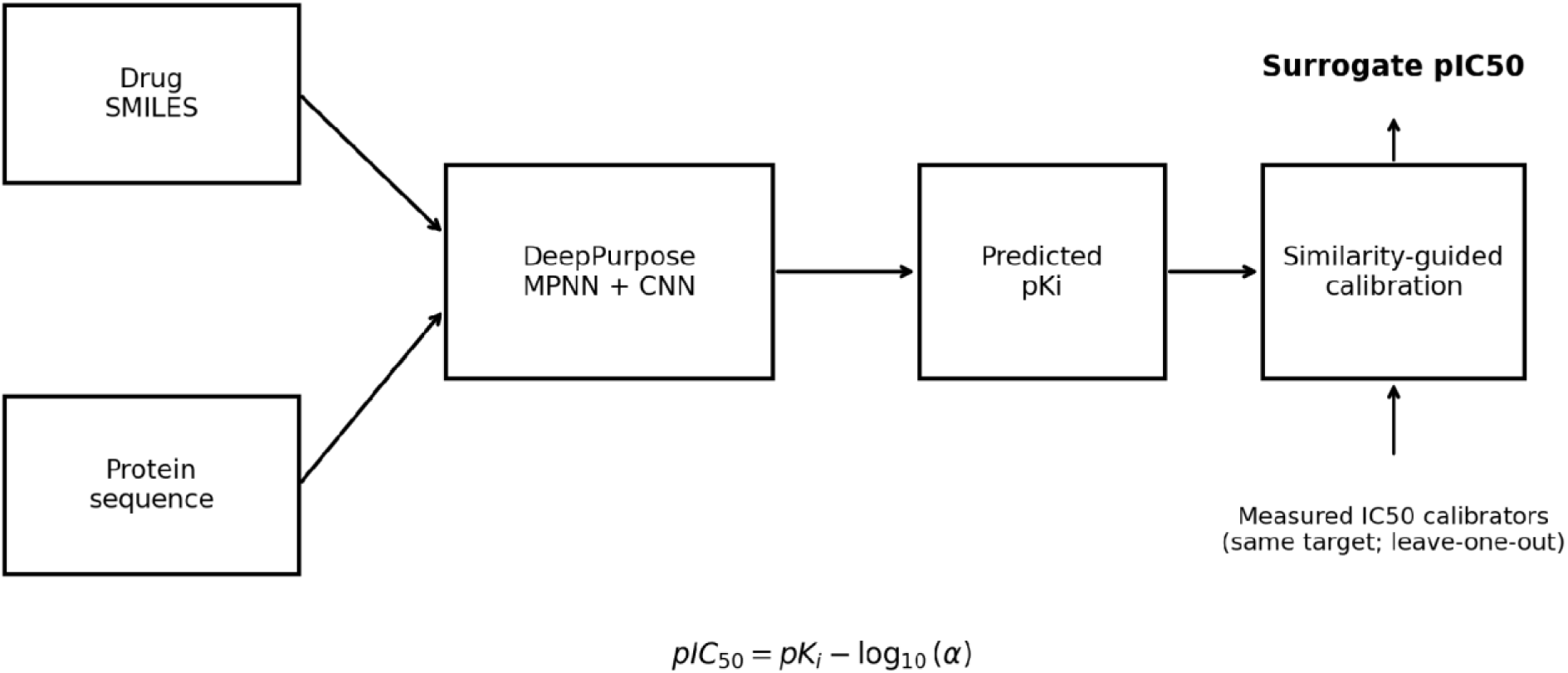
Two-stage workflow used in this study. DeepPurpose predicts pKi; a target-specific, similarity-guided empirical layer then estimates surrogate pIC50.

### 2.2 DeepPurpose architecture and training

Both checkpoints used the DeepPurpose framework.^1^ The compound encoder was a message-passing neural network (MPNN), a graph-neural-network model class introduced for molecular learning,^11^ configured here with three message-passing steps and a 128-dimensional hidden representation. The protein encoder was a one-dimensional convolutional neural network with 32, 64, and 96 filters and kernel widths of 4, 8, and 12. DeepPurpose supports molecular-graph message-passing encoders and protein-sequence CNN encoders and combines learned compound and protein representations in a regression decoder.^1^ In our configuration, the two representations were concatenated before a fully connected regression head trained with a squared-error objective. Batch size was 512, learning rate was 5 × 10−4, and the data split was 70/10/20. The original BindingDB and ChEMBL checkpoint was trained for 400 epochs.

#### 2.2.1 Long-protein sequence handling

The DeepPurpose implementation used here limits the protein-sequence input presented to the encoder to 1,000 amino acids. To avoid silent truncation of longer targets, we introduced an explicit tiered sequence-handling policy. Sequences of 1,000 residues or fewer were passed in full; sequences of 1,001–1,100 residues were truncated and flagged as minor truncations. For proteins longer than 1,100 residues, an annotated functional-domain span was used to determine whether the first 1,000 residues retained the domain or whether a domain-containing sequence window should instead be supplied. Targets longer than 1,100 residues without an available domain span were not predicted. For every accepted prediction, the sequence strategy and the sequence actually presented to the model were recorded in AffinityPrediction.metadata. Full decision rules and target-space prevalence data are provided in Supporting Information Section S11.1.

**Table 1.** Summary of the two DeepPurpose checkpoints. The all-organism ChEMBL build contained 339,023 eligible pairs; 150,000 were sampled for training because featurizing all unique molecules.

| Characteristic | BindingDB checkpoint | ChEMBL 37 checkpoint |
| --- | --- | --- |
| Training endpoint | Ki | Ki |
| Pairs used | 91,644 | 150,000 |
| Unique compounds represented | 39,342 | 112,881 |
| Unique targets represented | 4,172 | 1,824 |
| Assay restriction | Mixed / unspecified | ChEMBL Binding assays (B) |
| pKi 5.00 records | 23,015 (25.1%) | 0 |
| Epochs | 400 | 200 |

### 2.3 ChEMBL 37 training-set construction

ChEMBL 37 was queried directly.^4,7^ Eligible records had standard_type = Ki, standard_relation = ‘=’, assay_type = ‘B’, no ChEMBL data-validity comment, potential_duplicate = 0, target_type = ‘SINGLE PROTEIN’, and an available protein sequence. In ChEMBL, assay_type = ‘B’ denotes a Binding assay; it should not be interpreted as a guarantee that every record is cell-free or biochemical.^8^ Activity values were converted to nM and then to pKi = 9 − log10(Ki[nM]). Records at exactly pKi 5.00 were removed. Replicate measurements were collapsed by median pKi. Targets with insufficient data were excluded, and a replicate-noise filter removed targets with a per-target replicate standard deviation above 0.7 log units. Three extraction builds were recorded: a human-only build with at least 20 compounds per target (255,854 pairs, 601 targets), a human-only build with at least 3 compounds (259,201 pairs, 1,010 targets), and the final all-organism build with at least 3 compounds (339,023 pairs, 1,869 targets). The last build was the source for the ChEMBL checkpoint. To reduce confounding from differences in training-set size, the ChEMBL dataset was sampled to 150,000 compound–target pairs, yielding a training scale comparable to that of the original checkpoint while retaining the revised data-quality filters. This design allowed the effect of training-data curation to be examined without simultaneously introducing the full increase in dataset size. A separate biochemical-versus-cellular analysis was performed to assess assay-context heterogeneity and replicate variability; this analysis is reported in the Supporting Information and was not used to define the ChEMBL assay-type filter.

### 2.4 Evaluation datasets

Four evaluation settings were used. (i) The Davis kinase benchmark was used to assess cross-database transfer of the BindingDB checkpoint. Davis and co-workers originally profiled 72 inhibitors against 442 kinases.^9^ The DTA benchmark derived from these data was evaluated using the fixed test fold distributed with DeepDTA (5,010 pairs).^2,12^ The checkpoint received no Davis data during training. (ii) A kinase panel was used for the head-to-head comparison of the two checkpoints. Canonical SMILES strings were used for compound identity matching.^13^ Of the 2,000 records in the panel, 55 compounds (2.8%) also occurred in the ChEMBL 37 training set and were removed, leaving 1,945 records across 37 proteins. We refer to this set as the leakage-cleaned kinase panel. (iii) FLT3 and JAK2 were used as dense single-target sets. (iv) A non-kinase panel contained 14,823 compound–target pairs from 14 ChEMBL targets spanning GPCRs, proteases, enzymes, nuclear receptors, a transporter, and an ion channel.

### 2.5 Similarity-guided empirical calibration toward IC50

The calibration layer was defined on the logarithmic activity scale. For a given compound, the model first generated predicted pKi. Other compounds with measured IC50 against the same target were then used as calibrators. Each calibrator contributed the residual rj = predicted pKij − measured pIC50j. Morgan/extended-connectivity-style circular fingerprints (radius 2, 2,048 bits) were generated with RDKit.^14–16^ Tanimoto similarity was used to compare the query with each calibrator.^17^ Calibrators below threshold T were discarded, and the remaining residuals were combined by a similarity-weighted median. The held-out compound was never used to calibrate itself. The resulting correction is equivalent to estimating a multiplicative factor α in IC50 ≈ αKi, giving pIC50 = pKi − log10(α). This is an empirical mapping, not a mechanistic Cheng–Prusoff conversion.^5^

Similarity thresholds from 0.3 to 0.8 and minimum-calibrator counts from one to five were evaluated. If fewer than the required number of calibrators survived, the method abstained and the compound counted against coverage. The main summary therefore reports both accuracy and coverage.

### 2.6 Performance metrics

Absolute error was summarized using RMSE, MAE, bias, and fold error. For p-scale predictions, fold error for one observation was 10^|predicted − measured|. Median fold error was used as the main practical measure because, in these evaluations, arithmetic means were dominated by a small number of extreme errors. Ranking was evaluated using Pearson correlation, Spearman correlation, and concordance index (CI); CI is also used in DeepPurpose and established DTA benchmarks.^1,2^ For datasets larger than 5,000 rows, the CI implementation used fixed-seed subsampling, as recorded in the technical log.

### 2.7 Literature comparison

Literature values were not recomputed. DeepDTA and GraphDTA Davis results were taken from their published reports.^2,10^ These comparisons are descriptive rather than head-to-head: the published models were trained on Davis, whereas the BindingDB checkpoint evaluated here was applied to Davis without retraining.

## 3. Results and Discussion

### 3.1 The original DTA checkpoint transferred to an independent kinase benchmark

The BindingDB checkpoint reached a CI of 0.860 and an MSE of 0.382 on the Davis published test fold. DeepDTA reported CI 0.878 and MSE 0.261 for its 1D/1D model,^2^ while GraphDTA reported CI 0.893 and MSE 0.229 for its GIN variant.^10^ Those published models were trained on Davis, whereas our checkpoint was not. The ordering signal nevertheless transferred: the zero-shot CI was within 0.018 of DeepDTA and 0.033 of GraphDTA. This result supports use of the model for cross-target affinity estimation, but it does not establish equivalence to models trained directly on the benchmark.

**Figure 2.**
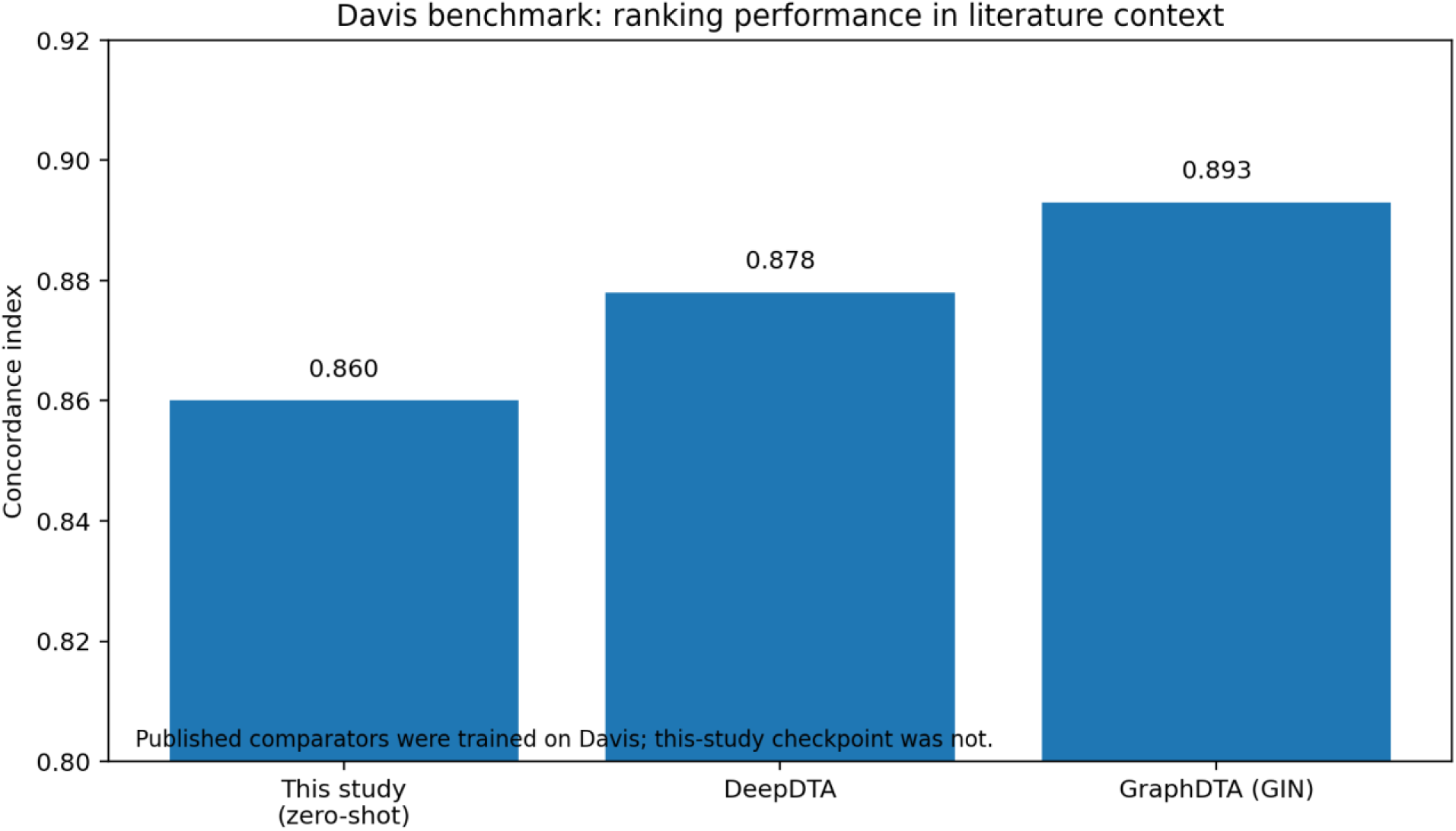
Davis benchmark concordance index in literature context. Published comparators were trained on Davis; the checkpoint from this study was evaluated zero-shot.

### 3.2 Cleaning the Ki training data reduced systematic error before calibration

The development data exposed a substantial problem in the original BindingDB training set: 23,015 of 91,644 labels (25.1%) were exactly pKi 5.00. The retrained ChEMBL set removed that censoring pattern and restricted training to exact Ki records from ChEMBL Binding assays. On the leakage-cleaned kinase panel, RMSE improved from 1.774 to 1.401, MAE from 1.427 to 1.132, CI from 0.510 to 0.577, and median fold error from 16.35× to 9.23×. The median bias shifted from −0.761 to +0.182 log units. Both R² values remained negative (−0.994 and −0.245), which is important: this 37-protein panel is close to a single-target regime and should not be presented as a strong ranking result.

The same trend was visible outside kinases. Across the 14-target non-kinase panel, the ChEMBL checkpoint reduced median fold error from 15.30× to 7.90×, mean RMSE from 1.701 to 1.328, and mean absolute bias from 0.856 to 0.441. CI improved on 13 of 14 targets, with the panel mean increasing from 0.522 to 0.600. The glucocorticoid receptor was the exception (CI 0.542 to 0.492), and HDAC1 remained a problematic target despite a small CI gain. These exceptions argue against presenting data cleaning as a universal cure; it improved the aggregate model, but target-specific failure modes remain.

**Figure 3.**
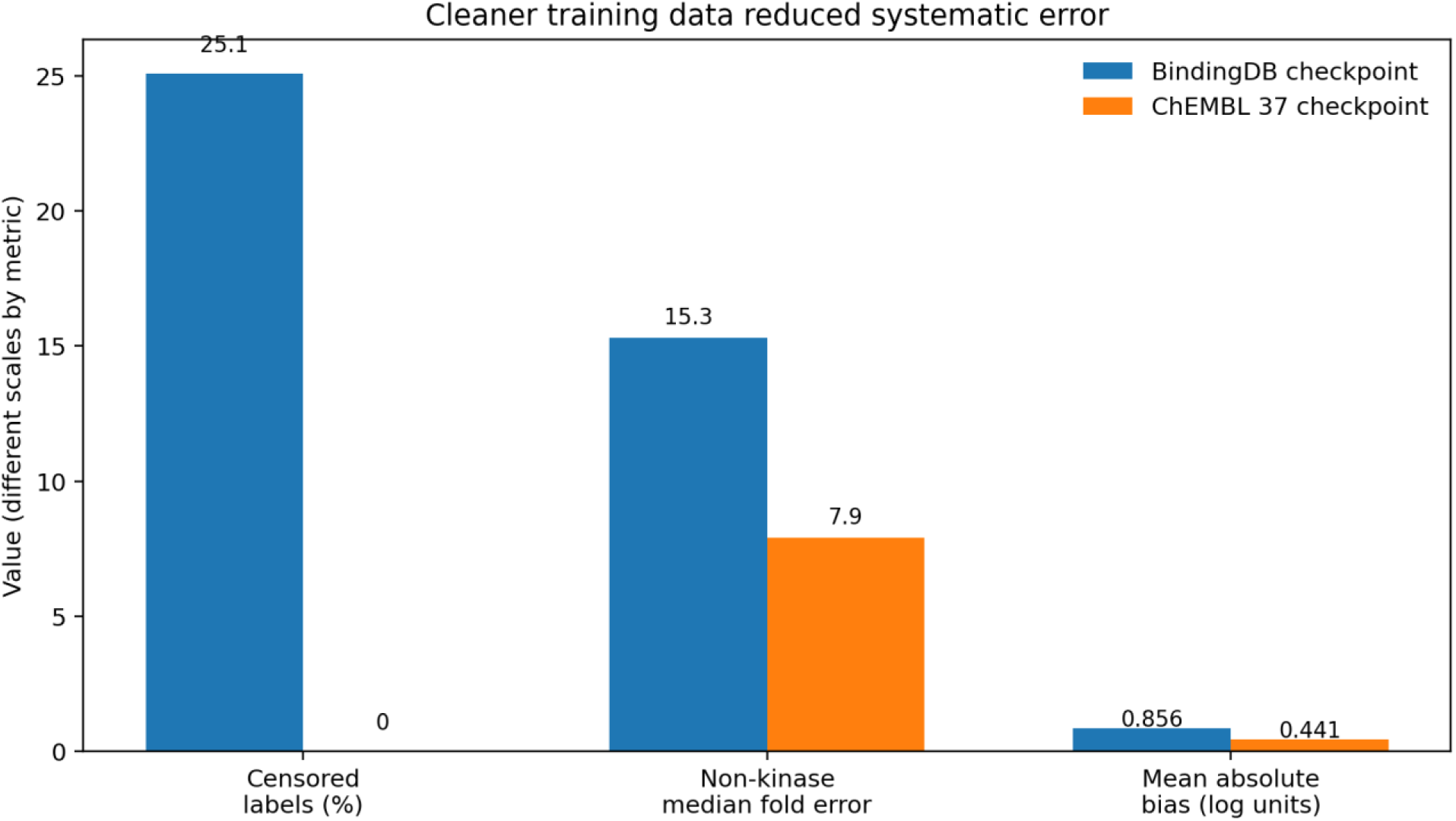
Effect of rebuilding the Ki training data. Values summarize censoring and non-kinase performance of the two checkpoints; metrics use different numerical scales and are shown together only to illustrate direction and magnitude of change.

**Table 2.** Head-to-head performance on the leakage-cleaned 1,945-record kinase panel.

| Metric | BindingDB checkpoint | ChEMBL 37 checkpoint |
| --- | --- | --- |
| RMSE | 1.774 | 1.401 |
| MAE | 1.427 | 1.132 |
| R<sup>2</sup> | −0.994 | −0.245 |
| Pearson | 0.022 | 0.261 |
| Spearman | 0.032 | 0.227 |
| Concordance index | 0.510 | 0.577 |
| Median fold error | 16.35× | 9.23× |
| Median bias | -0.761 | +0.182 |

### 3.3 Similarity-guided calibration aligned model output with measured IC50

The central question was whether predicted Ki could be made more useful for an IC50-based workflow without pretending that the two endpoints are identical. On the original non-kinase panel, using predicted pKi directly as pIC50 produced a median fold error of 15.30×. Similarity-guided calibration at T = 0.6 with one required calibrator reduced the panel median to 3.12× while retaining 85.7% coverage. For the ChEMBL checkpoint, the corresponding values were 7.90× before calibration and 3.22× after calibration. In other words, better Ki training roughly halved the uncalibrated error, but calibration still provided a further improvement of more than twofold in typical error.

The calibration was defined from the outset as empirical rather than mechanistic. The retraining experiment explains why that distinction matters. Once the affinity model was less biased, roughly half of the apparent correction disappeared before calibration was applied. The residual adjustment is therefore best described as a target- and chemistry-dependent offset between the model output and the observed IC50 assay space, not as a universal Ki-to-IC50 conversion factor.

The size of the remaining error should also be read against the reproducibility of the source measurements. In a separate project-level stratification of ChEMBL 37 records into biochemical and cellular subsets, the biochemical Ki subset had a median replicate SD of 0.382 log units (2.41×) and the biochemical IC50 subset 0.519 log units (3.30×). Because many replicate sets contained only three measurements, the Ki SD was biased downward; the recorded small-sample correction placed its underlying noise closer to 2.8×. Thus, a calibrated median error near 3.2× is on the same order as the replicate variability observed in that analysis. This does not establish a hard performance ceiling. The exact rule used to create the secondary biochemical-versus-cellular stratification was not preserved in the current technical record, so these values are used only as empirical context and should not be conflated with ChEMBL’s assay_type = ‘B’ definition.

**Table 3.** Non-kinase panel median fold error before and after calibration. Higher similarity thresholds reduce error at the cost of coverage.

| Setting | Coverage | BindingDB checkpoint | ChEMBL checkpoint |
| --- | --- | --- | --- |
| Uncalibrated | 100% | 15.30× | 7.90× |
| T = 0.4, min cal. = 1 | 95.9% | 3.55× | 3.50× |
| T = 0.6, min cal. = 1<br>(recommended) | 85.7% | 3.12× | 3.22× |
| T = 0.6, min cal. = 5 | 54.3% | 2.87× | 3.15× |
| T = 0.8, min cal. = 4 | 11.2% | 2.61× | 3.01× |

### 3.4 Similarity, rather than calibrator count, controlled the useful accuracy–coverage trade-off

A full threshold sweep on the BindingDB checkpoint showed a clear knee around T = 0.6. Raising T from 0.3 to 0.6 improved median fold error from 4.21× to 3.12× while coverage fell from 98.0% to 85.7%. Raising T further to 0.8 improved the median only to 2.89×, but coverage fell to 48.0%. In these data, higher similarity thresholds selected more accurate calibrations but left more queries unsupported.

**Figure 4.**
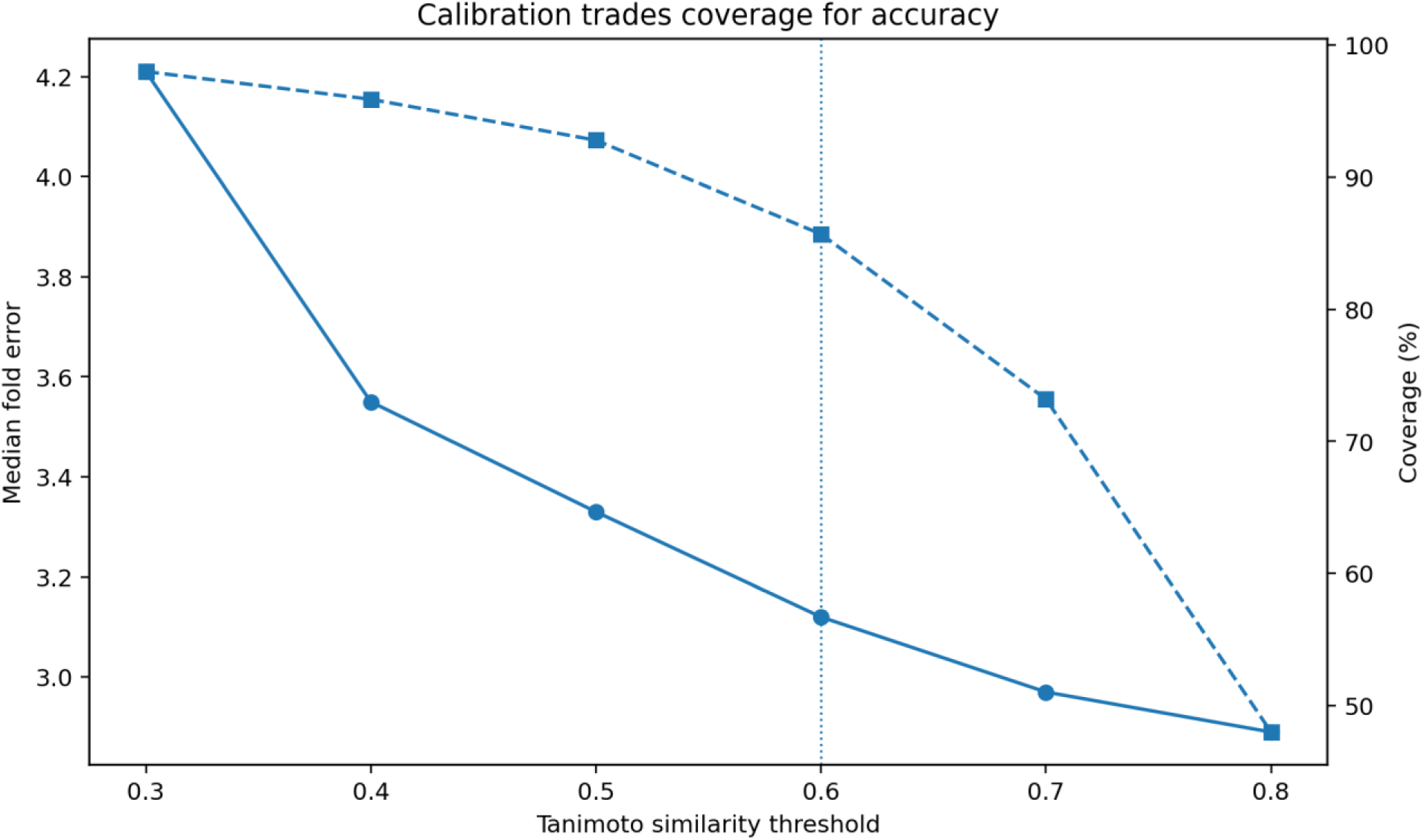
Accuracy–coverage trade-off across Tanimoto thresholds on the 14-target non-kinase calibration panel (BindingDB checkpoint; minimum calibrators = 1).

By contrast, the minimum number of calibrators had little effect on typical error. On FLT3 at T ≥ 0.6, compounds with exactly one available calibrator had a median fold error of 3.17×, compared with 3.12× for compounds with 10–24 or 25–49 calibrators. The decisive comparison was the group rejected by the old five-calibrator gate: 640 compounds with only one to four qualifying neighbors had a calibrated median error of 3.25×, almost identical to the 3.20× error among compounds with at least five neighbors; left uncalibrated, those same 640 compounds had 13.43× median error. Requiring five calibrators therefore removed useful predictions rather than isolating a clearly unreliable subset. The default was changed from five calibrators to one. This should not be read as evidence that one neighbor is always sufficient; it shows only that a count threshold adds little once a meaningful similarity threshold has already been imposed. The 50+ neighbor group was an exception, with 4.26× median error; more neighbors therefore did not guarantee a better local correction, and the reason for this degradation was not tested here.

**Figure 5.**
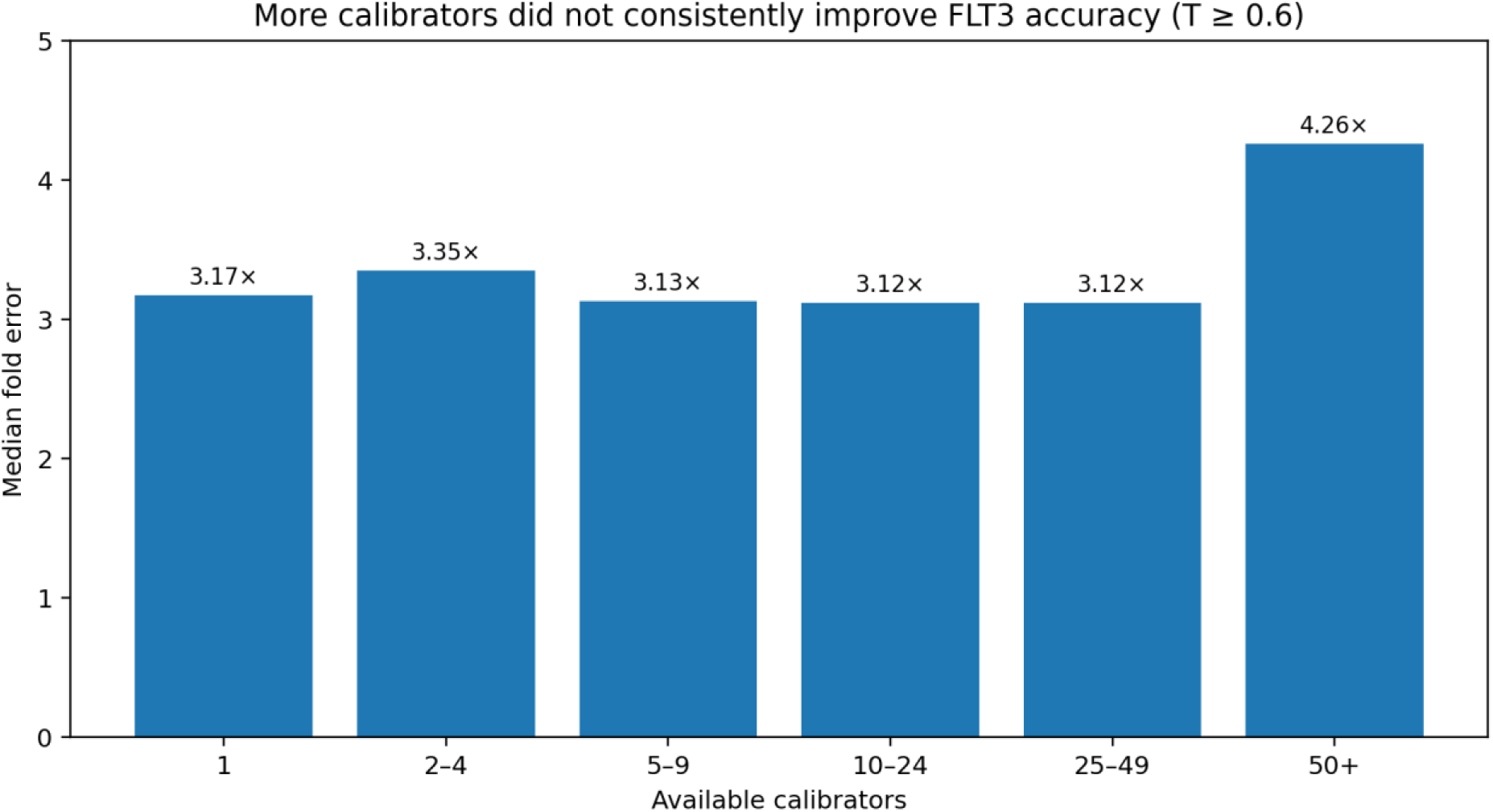
FLT3 median fold error by number of available calibrators at T ≥ 0.6. Accuracy was broadly flat from one to 49 neighbors.

### 3.5 The calibration behavior was not limited to kinases

The 14-target panel was assembled to test whether the empirical correction depended on kinase biology. It included four GPCRs, three proteases, three enzymes, two nuclear receptors, SERT and hERG. At T = 0.6 with one calibrator, the panel median error was 3.12× for the BindingDB checkpoint. Calibration improved every represented family, but the magnitude was not uniform: the median improvement was 11.45× for GPCRs, 7.61× for nuclear receptors, 4.32× for proteases, 3.87× for the ion channel, 3.46× for enzymes and 3.41× for the transporter. Across individual targets, improvement ranged from 2.75× for HDAC1 to 24.46×. The effect was therefore general across this panel, but clearly family- and target-dependent, **Tables 5 and S7**. The per-target analysis also shows why a single global factor is inadequate: the average correction required by the original checkpoint ranged from approximately 0.05× for thrombin to 45.53× for the D2 dopamine receptor, with the direction of the offset changing across families.

**Table 4.**
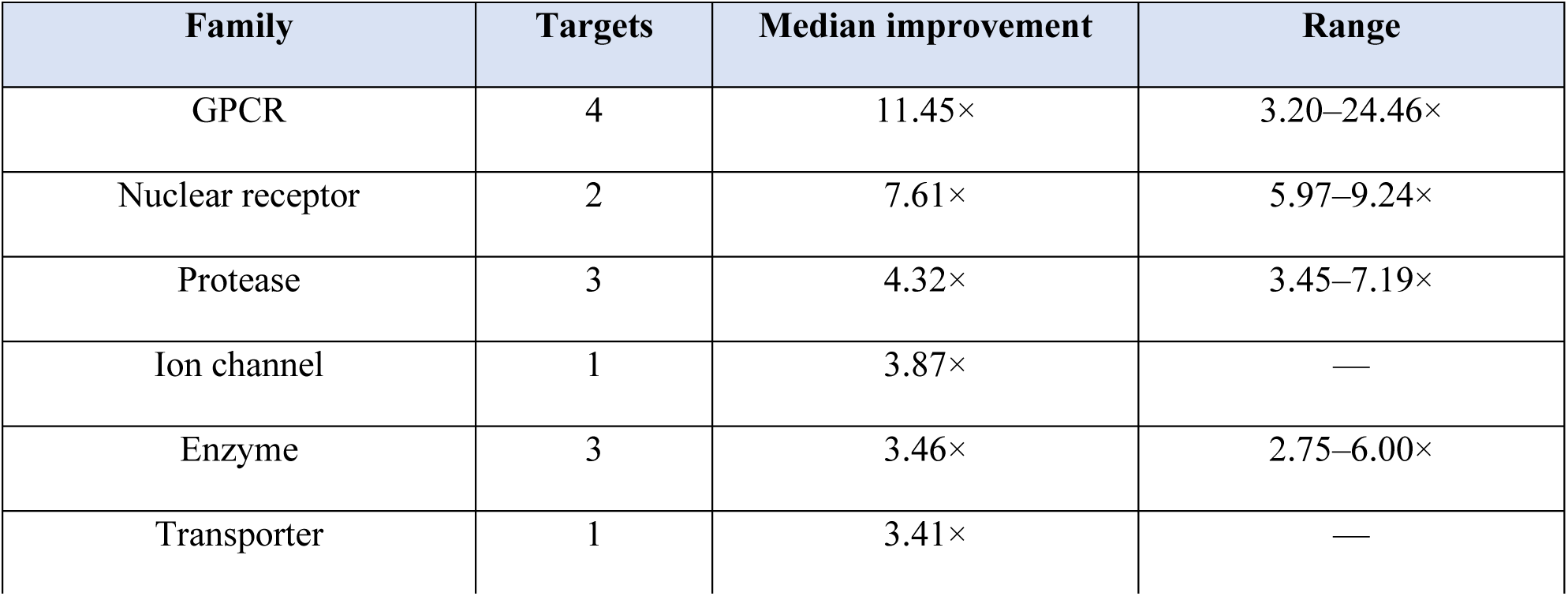
Calibration improvement by target family for the BindingDB checkpoint at T = 0.6 and minimum calibrators = 1. Improvement is the ratio of uncalibrated to calibrated median fold error.

**Figure 6.**
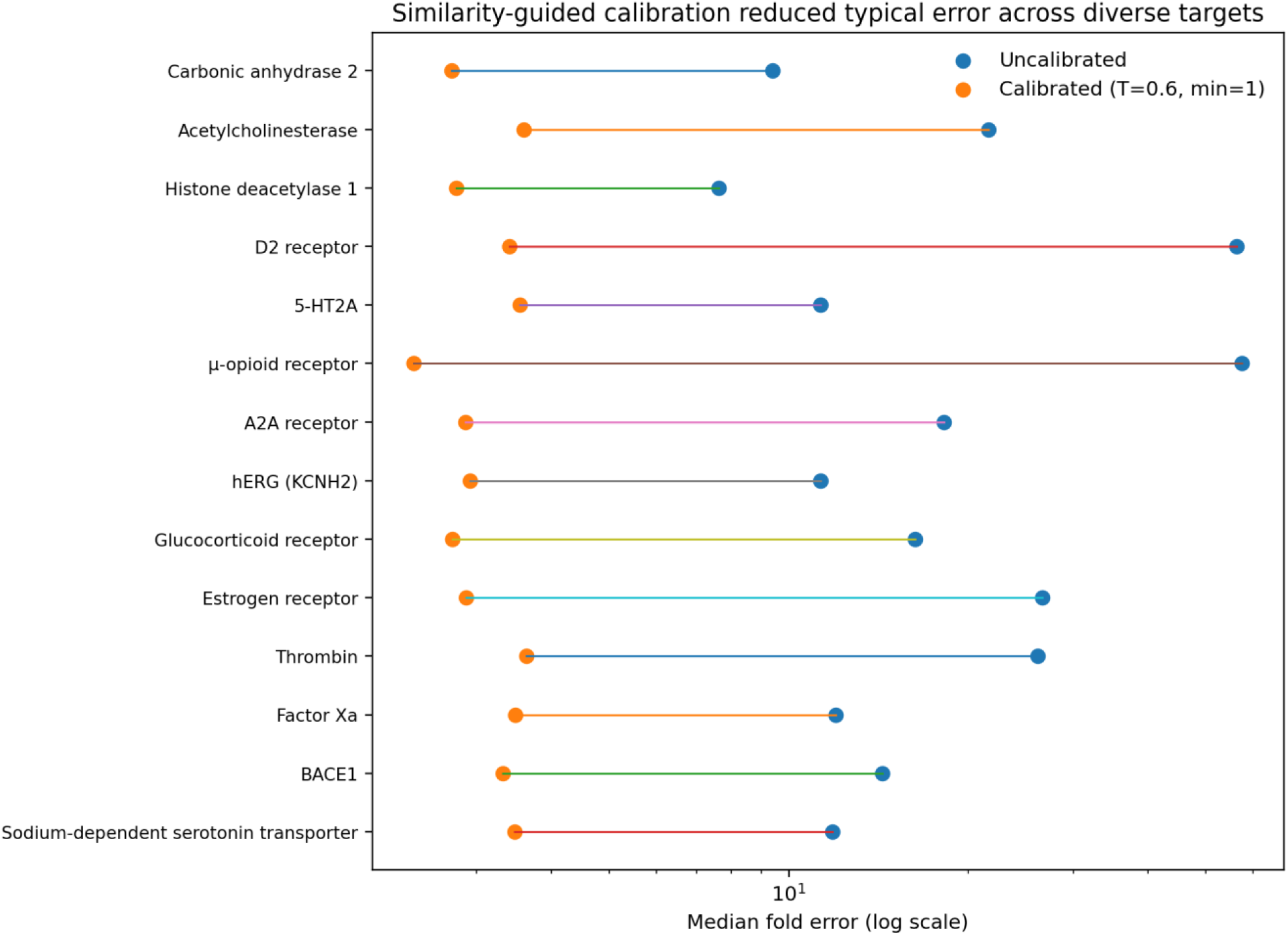
Per-target median fold error before and after similarity-guided calibration at T = 0.6 and minimum calibrators = 1. Values are from the 14-target non-kinase sweep using the BindingDB checkpoint.

### 3.6 Calibration improved absolute agreement but did not solve within-target ranking

The main limitation emerged when the protein was held fixed. The BindingDB checkpoint achieved CI 0.860 on the multi-target Davis set, but CI fell to 0.584 on FLT3 and 0.513 on JAK2. Spearman correlation similarly fell from 0.647 on Davis to 0.252 and 0.034. A single-assay FLT3 control did not rescue ranking (CI 0.562), which argues against pooled assay heterogeneity as the main explanation.

This behavior is consistent with the architecture used here. DeepPurpose concatenates learned compound and protein representations before the prediction decoder.^1^ When one target is fixed, the protein vector is constant for every compound and therefore cannot itself contribute to ordering within that target. The calibration layer cannot repair this because it rescales individual predictions; it does not add a target-specific ranking objective. The present system is therefore more defensible for cross-target triage and approximate endpoint alignment than for ranking a congeneric series against one protein.

### 3.7 Limitations and reproducibility considerations

Several limitations should remain visible in the final interpretation. First, both training runs used random splits rather than scaffold-disjoint or cold-target splits. Evaluation design is known to materially affect apparent drug– target prediction performance, especially when drugs or targets overlap between training and test data;^18^ the internal test metrics here should therefore not be interpreted as a formal test of extrapolation to unseen scaffolds or targets. Second, the BindingDB release used for the original checkpoint was not recorded, so that historical extract cannot be reproduced exactly. Third, the similarity threshold in the calibration layer is a coverage rule for access to local calibrators, not an applicability-domain estimate for the underlying DTA model. Fourth, high-affinity compounds remain difficult: the technical record reports a 36.6× median error above pKi 9 for the current service configuration. Finally, the surrogate IC50 is a model-derived estimate and should not be reported as an experimental equivalent.

The sequence-length constraint affected a non-negligible fraction of the target space. Among 11,053 single-protein targets with an available sequence, 1,381 (12.5%) exceeded 1,000 residues; among 8,752 targets carrying bioactivity data, 1,108 (12.7%) exceeded this limit. The issue was more common among protein kinases, where 145 of 663 targets (21.9%) exceeded 1,000 residues and 115 (17.3%) exceeded 1,100 residues. These observations motivated explicit sequence routing rather than unrecorded truncation. The routing policy makes the model input auditable, but it was not evaluated here as an independent accuracy-improvement intervention (Supporting Information Section S11.1).

The current implementation used DeepPurpose,^1^ RDKit 2026.03,^16^ scikit-learn,^19^ SciPy,^20^ and PyTorch.^21^ Training and inference checks used a single NVIDIA A10G. Exact package versions other than the recorded RDKit 2026.03 release family were not retained in the technical record and are therefore not inferred here. Because only the RDKit release family, not the exact patch version, was recorded, no version-specific RDKit DOI is claimed. These details, together with the calibration sweep and target-level results, are provided in the Supporting Information.

## 4. Conclusion

This study addresses a practical gap between drug–target affinity prediction and potency reporting. DeepPurpose was used to predict Ki from compound and protein information, and a second, similarity-guided layer was used to calibrate that prediction toward IC50. The two stages should remain conceptually separate: the first estimates binding affinity; the second empirically aligns the output with a different experimental endpoint.

The data support three conclusions. First, the DTA model retained useful cross-target ranking signal when transferred to the Davis benchmark without retraining. Second, cleaning the Ki training data materially reduced systematic error, showing that calibration should not be used to hide avoidable model bias. Third, local chemical similarity was the main determinant of calibration accuracy, while requiring many calibrators mainly reduced coverage. Across a diverse non-kinase panel, calibration brought typical error to roughly threefold for compounds with adequate local support. In the project’s separate assay-context analysis, this was the same order as the 2.41× and 3.30× replicate variability measured for the subsets classified as biochemical Ki and biochemical IC50, respectively; those values provide empirical context rather than a theoretical lower bound.

The method is not a universal Ki-to-IC50 conversion and it is not a replacement for experimental measurement. Its appropriate role is narrower: a transparent surrogate-potency layer for cross-target triage when the underlying DTA model and a local IC50 calibration neighborhood are available. Within-target ranking remains a limitation of the present architecture and will require target-specific modeling or retraining rather than additional scaling.

## Data availability

The analyses used public bioactivity resources including BindingDB,^3^ ChEMBL,^4,7^ and the Davis kinase benchmark.^9^ The Supporting Information supplied with this manuscript contains the model configuration, ChEMBL extraction rules and build definitions, the project-level assay-context and replicate analyses, evaluation-set definitions, calibration sweeps, target-level results, single-target controls, and reproducibility notes. Per-compound model predictions and the exact ChEMBL extraction script will be provided upon request.

## Supporting Information Available

Model configuration; ChEMBL extraction and data-quality rules; assay-context and replicate analyses; evaluation-set definitions; complete calibration sweeps; target-level results; FLT3 calibrator and assay-matched controls; single-target ranking; reproducibility and known-issue notes. Additional supporting material documents the long-protein sequence-routing policy and the prevalence of targets exceeding the 1,000-residue protein-input limit across ChEMBL target classes.

