## Supplement Figure for "From Predicted Ki to Surrogate IC50: Similarity-Guided Empirical Calibration of Drug–Target Affinity Predictions"

#### S1. Model configuration

The DTA implementation used the DeepPurpose framework[1] with a molecular message-passing encoder; MPNNs are described by Gilmer and co-workers.[2]

| Component | Configuration |
|---|---|
| Drug encoder | MPNN; 3 message-passing steps; 128-dimensional hidden state |
| Protein encoder | 1D CNN; filters 32/64/96; kernel widths 4/8/12 |
| Fusion | Concatenation before fully connected regression head |
| Objective | Squared error |
| Batch size | 512 |
| Learning rate | 5 × 10−4 |
| Split | 70/10/20; seed 42 |
| BindingDB checkpoint | 400 epochs |
| ChEMBL checkpoint | 200 epochs |

#### S2. ChEMBL 37 extraction and data-quality rules

The ChEMBL 37 SQLite database was queried directly.[3,4] The following inclusion criteria were recorded in the technical log: standard_type = 'Ki'; standard_relation = '='; assay_type = 'B'; data_validity_comment IS NULL; potential_duplicate = 0; target_type = 'SINGLE PROTEIN'; and sequence IS NOT NULL. ChEMBL defines assay_type = 'B' as Binding; this category is not synonymous with a guarantee of a cell-free biochemical assay.[5]

Activity values were converted to nM and pKi, records at exactly pKi 5.00 were removed, replicate measurements were collapsed by median pKi, and the target-noise filter was applied before training-set construction.

The intermediate counts came from separate extraction builds rather than from one monotonic filtering sequence. Build A was restricted to Homo sapiens and required at least 20 compounds per target (255,854 pairs, 601 targets). Build B was also human-only but required at least 3 compounds per target (259,201 pairs, 1,010 targets); the 259,623 figure in the development log is the pre-minimum-compound intermediate from this build. Build C dropped the organism restriction, retained the three-compound minimum, and yielded 339,023 pairs across 1,869 targets. Build C was the source pool for the ChEMBL checkpoint, from which 150,000 pairs were sampled for training.

| **Build** | **Organism filter** | **Minimum compounds** | **Pairs** | **Targets** |
|---|---|---|---|---|
| A | Homo sapiens | 20 | 255,854 | 601 |
| B | Homo sapiens | 3 | 259,201 | 1,010 |
| C (training source) | All organisms | 3 | 339,023 | 1,869 |

**S2.1 Replicate-noise filter**

| **Replicate analysis** | **Value** |
|---|---|
| Targets with replicate data | 408 |
| Targets with ≥5 repeated pairs | 167 |
| Median replicate SD before filtering | 0.418 log units (2.62×) |
| p90 replicate SD | 0.824 log units (6.67×) |
| Noise threshold | SD ≤ 0.7 log units |
| Targets removed | 31 |
| Median replicate SD after filtering | 0.358 log units (2.28×) |

### S3. Assay-type analysis and replicate reproducibility

A separate project analysis stratified public Ki and IC50 records into biochemical and cellular groups to examine assay-context heterogeneity. This secondary grouping is distinct from ChEMBL's assay_type = 'B' definition (Binding).[5] The current technical record preserves the resulting counts and summary statistics but not the exact query or rule used to assign records to the biochemical and cellular groups. The values below are therefore reported as a project-level sensitivity and reproducibility analysis, not as a definition supplied by ChEMBL. In total, 1,406,308 Ki and IC50 records across 3,098 targets were summarized, with 272 target/endpoint combinations supporting biochemical-versus-cellular comparisons. The direction of the shift was inconsistent: cellular measurements were weaker on 62% of IC50 comparisons and 66% of Ki comparisons, while target-level shifts ranged from 0.015× to 3,087×. The systematic component of the biochemical-to-cellular shift was 0.26–0.52 log units against a per-target spread of 0.74–1.08 log units (signal-to-noise ratio 0.35–0.48). Within this project-level stratification, assay context was therefore not the dominant source of heterogeneity.

The training extraction itself used ChEMBL Binding assays (assay_type = 'B'), not the secondary biochemical/cellular category. In the separate stratification, the subset classified as biochemical Ki had the lowest recorded replicate SD: 0.382 log units (2.41×), compared with 0.700 log units (5.01×) for the cellular Ki subset. The biochemical and cellular IC50 subsets had similar replicate SDs of 0.519 and 0.511 log units, respectively (3.30× and 3.24×). These measurements are used as empirical context for data variability, not as a justification for equating ChEMBL's Binding code with 'biochemical.'

| Endpoint | Project-level assay-context group | Records / replicate pairs | Median / geomean | Replicate SD | 1-SD fold |
|---|---|---|---|---|---|
| IC50 | Biochemical | 975,890 records; 20,451 replicate pairs | Median 143.0 nM | 0.519 | 3.30× |
| IC50 | Cellular | 67,367 records; 808 replicate pairs | Median 614.0 nM | 0.511 | 3.24× |
| Ki | Biochemical | 317,705 records; 4,393 replicate pairs | Median 68.9 nM | 0.382 | 2.41× |

| Endpoint | Project-level assay-context group | Records / replicate pairs | Median / geomean | Replicate SD | 1-SD fold |
|---|---|---|---|---|---|
| Ki | Cellular | 43,589 records; 77 replicate pairs | Median 631.0 nM | 0.700 | 5.01× |

For the project-defined biochemical versus cellular Ki replicate comparison, the recorded Mann–Whitney p value was 2.9 × 10−11. About 60% of repeated compound–target pairs had exactly three measurements; simulation of the n = 3 sample-SD bias gave a measured/true SD ratio of 0.83–0.84, placing the underlying variability of the project-defined biochemical Ki subset nearer 2.8× than the uncorrected 2.41× estimate. This correction is used only as context for the noise scale and not to alter any model metric.

**S4. Evaluation sets**

The original affinity checkpoint used BindingDB,[6] while the single-target and non-kinase evaluation sets were extracted from ChEMBL.[3] Davis and co-workers reported the original kinase selectivity panel,[7] and the fixed benchmark split follows DeepDTA and its distributed train/test folds.[8,9] Canonical SMILES strings were used for the ChEMBL-training overlap check.[10] Fifty-five compounds from the original 2,000-record kinase panel (2.8%) occurred in the ChEMBL 37 training set and were removed, leaving 1,945 records across 37 proteins.

| Dataset | Records / compounds | Targets | Purpose |
|---|---|---|---|
| Davis published test fold | 5,010 pairs | 379 in recorded fold | Cross-database transfer |
| Leakage-cleaned kinase panel | 1,945 records | 37 | Old vs new checkpoint |
| FLT3 (CHEMBL1974) | 3,743 compounds | 1 | Single-target ranking and calibration |
| JAK2 (CHEMBL2971) | 9,677 compounds | 1 | Single-target ranking and calibration |

| Dataset | Records / compounds | Targets | Purpose |
|---|---|---|---|
| Non-kinase panel | 14,823 pairs | 14 | Cross-family calibration |

**S5. Full non-kinase calibration sweep**

Calibration similarity was computed from RDKit-generated Morgan/extended-connectivity-style circular fingerprints[11–13] using the Tanimoto index.[14]

| T | Min cal. | Coverage | Median fold | Geo-mean fold | Within 3× |
|---|---|---|---|---|---|
| none | — | 100.0% | 15.30× | 23.33× | 20.9% |
| 0.3 | 1 | 98.0% | 4.21× | 5.91× | 39.8% |
| 0.3 | 2 | 96.6% | 4.20× | 5.85× | 39.9% |
| 0.3 | 3 | 95.2% | 4.22× | 5.91× | 39.8% |
| 0.3 | 4 | 92.8% | 4.25× | 5.98× | 39.5% |
| 0.3 | 5 | 90.7% | 4.23× | 5.90× | 39.7% |
| 0.4 | 1 | 95.9% | 3.55× | 5.07× | 43.8% |
| 0.4 | 2 | 92.3% | 3.53× | 5.03× | 44.2% |
| 0.4 | 3 | 89.4% | 3.49× | 4.97× | 44.6% |
| 0.4 | 4 | 86.5% | 3.48× | 4.94× | 44.5% |
| 0.4 | 5 | 83.8% | 3.50× | 4.95× | 44.5% |
| 0.5 | 1 | 92.8% | 3.33× | 4.78× | 46.2% |
| 0.5 | 2 | 88.1% | 3.28× | 4.74× | 47.2% |
| 0.5 | 3 | 82.7% | 3.21× | 4.61× | 47.8% |
| 0.5 | 4 | 76.9% | 3.27× | 4.53× | 47.2% |
| 0.5 | 5 | 71.7% | 3.24× | 4.53× | 47.8% |
| 0.6 | 1 | 85.7% | 3.12× | 4.48× | 48.6% |
| 0.6 | 2 | 75.5% | 2.97× | 4.24× | 50.3% |

| T | Min cal. | Coverage | Median fold | Geo-mean fold | Within 3× |
|---|---|---|---|---|---|
| 0.6 | 3 | 68.3% | 2.95× | 4.19× | 50.6% |
| 0.6 | 4 | 60.6% | 2.92× | 4.12× | 50.8% |
| 0.6 | 5 | 54.3% | 2.87× | 3.95× | 51.9% |
| 0.7 | 1 | 73.2% | 2.97× | 4.24× | 50.1% |
| 0.7 | 2 | 57.2% | 2.90× | 4.13× | 51.7% |
| 0.7 | 3 | 46.9% | 2.76× | 3.77× | 53.6% |
| 0.7 | 4 | 38.6% | 2.70× | 3.60× | 55.4% |
| 0.7 | 5 | 31.8% | 2.74× | 3.68× | 53.8% |
| 0.8 | 1 | 48.0% | 2.89× | 4.26× | 51.8% |
| 0.8 | 2 | 29.0% | 2.88× | 4.00× | 51.8% |
| 0.8 | 3 | 19.0% | 2.75× | 3.56× | 55.2% |
| 0.8 | 4 | 11.2% | 2.61× | 3.69× | 55.3% |
| 0.8 | 5 | 6.7% | 2.72× | 3.60× | 54.7% |

The preselected practical setting was T = 0.6 with minimum calibrators = 1. At that point the BindingDB checkpoint retained 85.7% coverage with a panel median fold error of 3.12×. More stringent thresholds continued to improve median error, but coverage fell sharply.

**S6. Target-level non-kinase results at T = 0.6, minimum calibrators = 1**

| Target | n | Uncalibrated median fold | Calibrated median fold | Coverage |
|---|---|---|---|---|
| Carbonic anhydrase 2 | 817 | 9.39× | 2.72× | 82.3% |
| Acetylcholinesterase | 1,200 | 21.63× | 3.60× | 85.3% |
| Histone deacetylase 1 | 1,200 | 7.64× | 2.77× | 84.9% |
| D2 receptor | 491 | 56.33× | 3.40× | 80.9% |
| 5-HT2A | 911 | 11.31× | 3.54× | 85.4% |
| μ-opioid receptor | 835 | 57.53× | 2.35× | 89.3% |

| Target | n | Uncalibrated median fold | Calibrated median fold | Coverage |
|---|---|---|---|---|
| A2A receptor | 969 | 18.18× | 2.87× | 94.4% |
| hERG (KCNH2) | 1,200 | 11.31× | 2.92× | 81.8% |
| Glucocorticoid receptor | 1,200 | 16.28× | 2.73× | 94.9% |
| Estrogen receptor | 1,200 | 26.61× | 2.88× | 77.6% |
| Thrombin | 1,200 | 26.11× | 3.63× | 91.3% |
| Factor Xa | 1,200 | 11.99× | 3.48× | 94.9% |
| BACE1 | 1,200 | 14.33× | 3.32× | 90.9% |
| Sodium-dependent serotonin transporter | 1,200 | 11.83× | 3.47× | 85.9% |

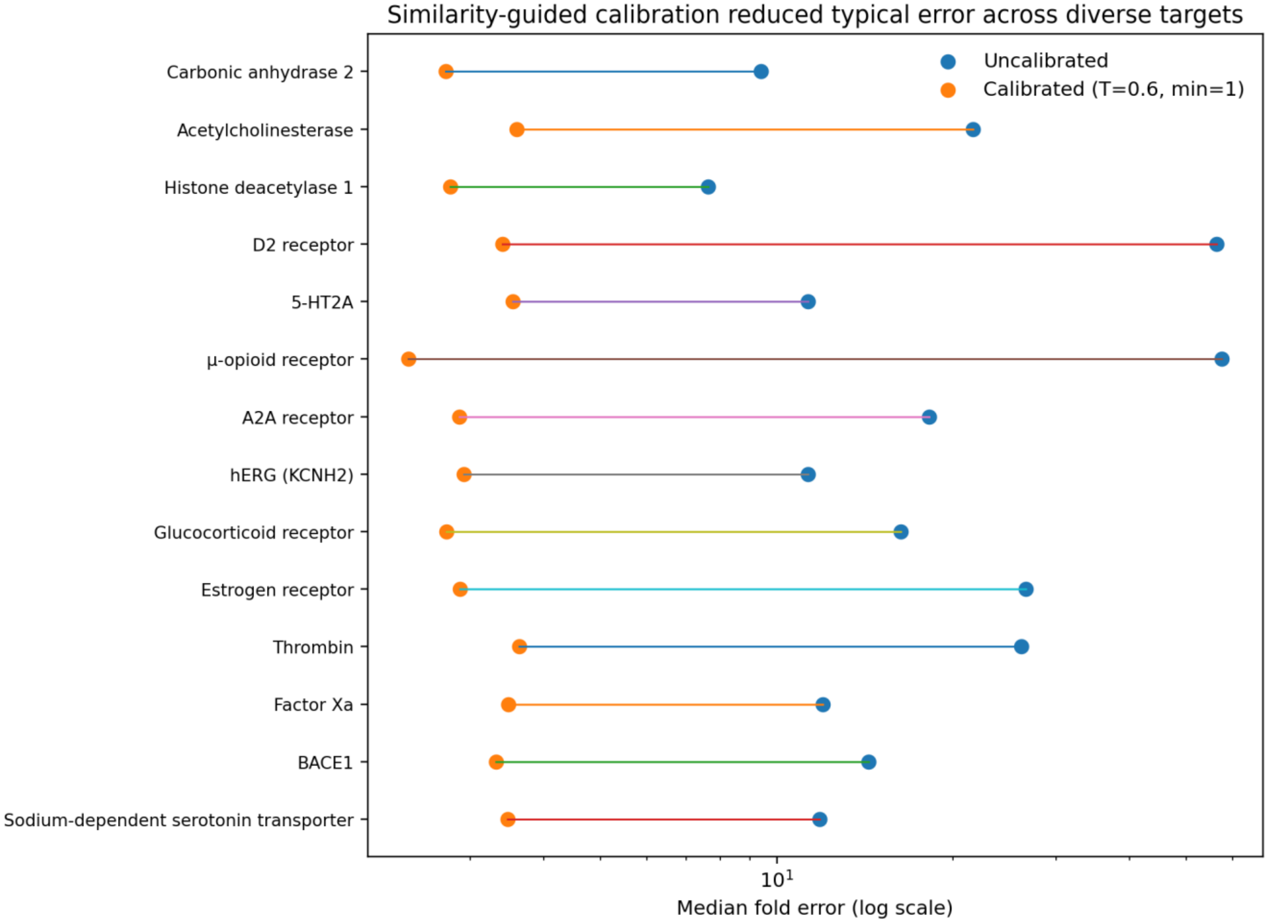


**Figure S1. Target-level change in median fold error at the T = 0.6, minimum-calibrator = 1 setting.**

**S7. FLT3 calibrator-count analysis**

| Available calibrators | n | Median fold error |
|---|---|---|
| Exactly 1 | 173 | 3.17× |
| 2–4 | 467 | 3.35× |
| 5–9 | 685 | 3.13× |
| 10–24 | 1,410 | 3.12× |
| 25–49 | 567 | 3.12× |
| 50+ | 196 | 4.26× |

At T ≥ 0.6, median error was essentially flat from one to 49 available calibrators. The 50+ group was worse (4.26×), showing that a larger calibrator pool did not guarantee better accuracy in this analysis. The cause of that degradation was not determined here.

**S8. FLT3 threshold and coverage analysis**

| Threshold | Coverage | Median fold error |
|---|---|---|
| None | 100% | 8.49× |
| T ≥ 0.4 | 97.2% | 4.02× |
| T ≥ 0.6 | 93.5% | 3.21× |
| T ≥ 0.8 | 58.5% | 2.73× |

The five-calibrator gate reduced FLT3 coverage to 76.4% at 3.20× median error. Reducing the gate to one calibrator increased coverage to 93.5% at 3.21×. More directly, the 640 compounds with only one to four qualifying calibrators had 3.25× median error after calibration versus 13.43× for those same compounds without calibration; the 2,858 compounds with five or more calibrators had 3.20× median error. The old gate was therefore excluding a group that calibrated nearly as well as the retained group. This is the basis for the current minimum-calibrator default of one.

| FLT3 group at T ≥ 0.6 | n | Median fold error |
|---|---|---|
| 5+ calibrators (retained by old gate) | 2,858 | 3.20× |

| FLT3 group at T ≥ 0.6 | n | Median fold error |
|---|---|---|
| 1–4 calibrators (excluded by old gate) | 640 | 3.25× |
| Same 1–4 group, uncalibrated | 640 | 13.43× |

#### S9. Assay-matched FLT3 control

| Calibrator pool | No similarity weighting | T ≥ 0.4 | T ≥ 0.6 |
|---|---|---|---|
| Mixed assay (3,743) | 7.81× | 5.59× | 5.07× |
| Same assay (273) | 5.63× | 5.58× | 4.95× |

Restricting calibrators to a single biochemical protocol improved the unweighted result, but at T ≥ 0.6 the mixed- and same-assay results were similar. On the same 273 compounds, single-target ranking did not improve (CI 0.562 versus 0.584 on the full FLT3 set).

#### S10. Single-target ranking

| Set | Distinct proteins | CI | Spearman |
|---|---|---|---|
| Davis | 379 | 0.860 | 0.647 |
| FLT3 | 1 | 0.584 | 0.252 |
| JAK2 | 1 | 0.513 | 0.034 |

Calibration did not change CI in the recorded tests because the α layer changes absolute values but does not introduce a new within-target ranking model.

#### S11. Reproducibility and known issues

Training used random 70/10/20 splits rather than scaffold-disjoint or cold-target splits; evaluation setting is known to materially affect apparent drug–target prediction performance.[15] Software dependencies were DeepPurpose,[1] RDKit,[13] scikit-learn,[16] SciPy,[17] and PyTorch.[18] The technical record retained only the RDKit 2026.03 release family; exact package versions for DeepPurpose, scikit-learn, SciPy, and PyTorch were not recorded and are not inferred in this SI. Because the exact RDKit patch version was not retained, no version-specific RDKit DOI is asserted.

| Item | Recorded status |
|---|---|
| Recorded software | DeepPurpose; RDKit 2026.03 (patch version not recorded); scikit-learn; SciPy; PyTorch |
| Hardware | Single NVIDIA A10G (23 GB); 30 GB system RAM |
| Random seed | 42 |
| Golden prediction check | Independent repeated outputs agreed to 3.0 × 10−6 pKi |
| Historical BindingDB release | Not recorded; exact historical extract is not reproducible |
| Original split handling | Training script concatenated files and re-split; claims about the on-disk held-out file require seed-42 verification |
| Applicability domain | Not defined; Tanimoto threshold applies only to calibration support |
| Service-layer σ = 0.8 pKi | Placeholder; must not be reported as calibrated uncertainty |
| Split type | Random; not scaffold-disjoint or cold-target |
| High-affinity limitation | Technical record reports 36.6× median error above pKi 9 |

#### S11.1 Long-protein sequence handling and target-space prevalence

The protein-sequence encoder in the DeepPurpose implementation used for this study receives at most 1,000 amino acids. Because unrecorded truncation can remove target-domain information, sequence handling was made explicit in the prediction service. The routing policy records what the model actually received. Domain spans were supplied by the upstream target annotation when available; this analysis does not claim that domain extraction itself improves predictive accuracy.

| Sequence length / condition | Model input | sequence_strategy |
|---|---|---|
| ≤ 1,000 aa | Whole sequence | full |
| 1,001–1,100 aa | First 1,000 residues; truncation explicitly flagged | minor_truncation |
| > 1,100 aa; domain ends ≤ 1,000 | First 1,000 residues; domain retained | truncated_domain_intact |

| | | |
|---|---|---|
| > 1,100 aa; domain extends beyond 1,000 | Domain-containing window of ≤ 1,000 residues | domain_extracted |
| > 1,100 aa; no domain span supplied | No prediction returned | rejected |

**Table S12. Sequence-routing policy for proteins approaching or exceeding the 1,000-residue model-input limit.**

Across the target inventory, 11,053 single-protein targets had an available sequence. Of these, 1,381 (12.5%) were longer than 1,000 residues and 1,090 (9.9%) were longer than 1,100 residues. The 1,001–1,100-residue tier contained 291 targets (2.6% of all sequence-bearing targets). Among 8,752 targets with bioactivity data, 1,108 (12.7%) exceeded 1,000 residues.

| Measure | Observed value |
|---|---|
| Single-protein targets with sequence | 11,053 |
| >1,000 aa | 1,381 (12.5%) |
| >1,100 aa | 1,090 (9.9%) |
| 1,001–1,100 aa | 291 (2.6%) |
| Median sequence length | 465 aa |
| Longest sequence | 34,350 aa |
| Targets with bioactivity data | 8,752 |
| >1,000 aa among bioactivity targets | 1,108 (12.7%) |

**Table S13. Overall prevalence of sequences exceeding the DeepPurpose input limit.**

The prevalence of long proteins varied substantially across broad target classes.

| Broad class | Targets | >1,000 aa n (%) | >1,100 aa n (%) | Median aa | Max aa |
|---|---|---|---|---|---|
| Epigenetic | 203 | 76 (37.4%) | 66 (32.5%) | 747 | 5537 |

| | | | | | |
|---|---|---|---|---|---|
| Ion | 437 | 120 (27.5%) | 115 (26.3%) | 640 | 5317 |
| Structural | 80 | 19 (23.8%) | 19 (23.8%) | 489 | 7393 |
| Adhesion | 42 | 7 (16.7%) | 7 (16.7%) | 675 | 2813 |
| Nuclear | 55 | 9 (16.4%) | 7 (12.7%) | 425 | 1621 |
| Unclassified | 2156 | 347 (16.1%) | 276 (12.8%) | 445 | 14507 |
| Transporter | 409 | 65 (15.9%) | 48 (11.7%) | 602 | 2261 |
| Auxiliary | 32 | 5 (15.6%) | 3 (9.4%) | 160 | 1150 |
| Enzyme | 5609 | 632 (11.3%) | 480 (8.6%) | 482 | 34350 |
| Cytosolic | 266 | 21 (7.9%) | 17 (6.4%) | 317 | 2843 |
| Membrane | 1137 | 57 (5.0%) | 32 (2.8%) | 386 | 2555 |
| Transcription | 354 | 15 (4.2%) | 12 (3.4%) | 468 | 2718 |
| Secreted | 214 | 8 (3.7%) | 8 (3.7%) | 228 | 5654 |
| Surface | 59 | 0 (0.0%) | 0 (0.0%) | 404 | 982 |

**Table S14. Sequence-length prevalence by broad ChEMBL target class.**

The most affected families are shown below (families with at least 10 targets).

| Family | Targets | >1,000 aa n (%) | >1,100 aa n (%) | Median aa | Max aa |
|---|---|---|---|---|---|
| NTPase ATP | 46 | 38 (82.6%) | 38 (82.6%) | 1426 | 2261 |
| NTPase P-type | 18 | 12 (66.7%) | 0 (0.0%) | 1021 | 1044 |
| Receptor toll-like | 22 | 10 (45.5%) | 0 (0.0%) | 904 | 1050 |
| Regulator writer | 42 | 18 (42.9%) | 18 (42.9%) | 834 | 3969 |
| Channel VGC | 209 | 85 (40.7%) | 83 (39.7%) | 889 | 4303 |

| | | | | | |
|---|---|---|---|---|---|
| Regulator eraser | 52 | 20 (38.5%) | 15 (28.8%) | 596 | 2540 |
| Regulator reader | 109 | 38 (34.9%) | 33 (30.3%) | 747 | 5537 |
| Transporter | 28 | 8 (28.6%) | 6 (21.4%) | 634 | 1521 |
| Receptor 7TM3 | 40 | 11 (27.5%) | 7 (17.5%) | 912 | 1215 |
| Structural | 80 | 19 (23.8%) | 19 (23.8%) | 489 | 7393 |
| Kinase protein | 663 | 145 (21.9%) | 115 (17.3%) | 628 | 4128 |
| Channel LGIC | 164 | 30 (18.3%) | 29 (17.7%) | 511 | 5317 |
| Phosphatase protein | 107 | 18 (16.8%) | 14 (13.1%) | 473 | 2485 |
| Adhesion | 42 | 7 (16.7%) | 7 (16.7%) | 675 | 2813 |
| Unclassified | 2156 | 347 (16.1%) | 276 (12.8%) | 445 | 14507 |
| Enzyme | 411 | 66 (16.1%) | 55 (13.4%) | 515 | 7073 |
| Transport protein | 32 | 5 (15.6%) | 3 (9.4%) | 160 | 1150 |
| Transferase | 1131 | 168 (14.9%) | 133 (11.8%) | 435 | 34350 |
| Receptor | 169 | 24 (14.2%) | 16 (9.5%) | 427 | 2555 |
| Ligase | 182 | 23 (12.6%) | 13 (7.1%) | 583 | 2458 |

**Table S15. Most affected target families.**

Several families were minimally affected or fully below the limit.

| **Family** | **Targets** | **>1,000 aa n (%)** | **>1,100 aa n (%)** | **Median aa** | **Max aa** |
|---|---|---|---|---|---|

| | | | | | |
|---|---|---|---|---|---|
| Electrochemical SLC | 262 | 5 (1.9%) | 3 (1.1%) | 580 | 1212 |
| Receptor 7TM1 | 785 | 1 (0.1%) | 1 (0.1%) | 374 | 1396 |
| Antigen | 59 | 0 (0.0%) | 0 (0.0%) | 404 | 982 |
| Cytochrome P450 | 103 | 0 (0.0%) | 0 (0.0%) | 503 | 561 |
| Protease aspartic | 48 | 0 (0.0%) | 0 (0.0%) | 409 | 876 |
| Factor nuclear | 126 | 0 (0.0%) | 0 (0.0%) | 470 | 984 |
| Receptor 7TMTAS2R | 25 | 0 (0.0%) | 0 (0.0%) | 309 | 338 |
| Receptor 7TMFZ | 16 | 0 (0.0%) | 0 (0.0%) | 616 | 793 |
| NTPase F-type | 15 | 0 (0.0%) | 0 (0.0%) | 511 | 838 |
| Phosphodiesterase PDE4 | 12 | 0 (0.0%) | 0 (0.0%) | 736 | 886 |
| Protease threonine | 19 | 0 (0.0%) | 0 (0.0%) | 241 | 420 |
| Phosphodiesterase | 16 | 0 (0.0%) | 0 (0.0%) | 449 | 888 |

**Table S16. Least affected target families.**

Among 5,609 enzyme targets, 632 (11.3%) exceeded 1,000 residues; protein kinases were particularly affected.

| Enzyme family | Targets | >1,000 aa n (%) | >1,100 aa n (%) | Median aa | Max aa |
|---|---|---|---|---|---|
| Lyase guanylate | 6 | 5 (83.3%) | 0 (0.0%) | 1052 | 1073 |
| Kinase protein | 663 | 145 (21.9%) | 115 (17.3%) | 628 | 4128 |

| | | | | | |
|---|---|---|---|---|---|
| Phosphatase protein | 107 | 18 (16.8%) | 14 (13.1%) | 473 | 2485 |
| Enzyme | 411 | 66 (16.1%) | 55 (13.4%) | 515 | 7073 |
| Transferase | 1131 | 168 (14.9%) | 133 (11.8%) | 435 | 34350 |
| Ligase | 182 | 23 (12.6%) | 13 (7.1%) | 583 | 2458 |
| Kinase | 8 | 1 (12.5%) | 1 (12.5%) | 440 | 1102 |
| Kinase regulator | 25 | 3 (12.0%) | 2 (8.0%) | 381 | 1332 |
| Phosphatase | 43 | 5 (11.6%) | 5 (11.6%) | 524 | 1445 |
| Protease cysteine | 114 | 13 (11.4%) | 11 (9.6%) | 427 | 3391 |
| Protease metallo | 150 | 14 (9.3%) | 11 (7.3%) | 647 | 1427 |
| Hydrolase | 1215 | 100 (8.2%) | 64 (5.3%) | 462 | 2897 |
| Lyase | 189 | 12 (6.3%) | 9 (4.8%) | 336 | 1610 |
| Isomerase | 145 | 8 (5.5%) | 8 (5.5%) | 384 | 1626 |
| Reductase | 767 | 40 (5.2%) | 31 (4.0%) | 418 | 2540 |
| Protease serine | 198 | 7 (3.5%) | 5 (2.5%) | 463 | 1860 |
| Protease | 6 | 0 (0.0%) | 0 (0.0%) | 786 | 844 |
| Protease threonine | 19 | 0 (0.0%) | 0 (0.0%) | 241 | 420 |
| Protease aspartic | 48 | 0 (0.0%) | 0 (0.0%) | 409 | 876 |
| Phosphodiesterase PDE4 | 12 | 0 (0.0%) | 0 (0.0%) | 736 | 886 |
| Phosphodiesterase PDE7 | 5 | 0 (0.0%) | 0 (0.0%) | 450 | 482 |
| Phosphodiesterase PDE6 | 7 | 0 (0.0%) | 0 (0.0%) | 858 | 861 |

| | | | | | |
|---|---|---|---|---|---|
| Phosphodiesterase PDE1 | 5 | 0 (0.0%) | 0 (0.0%) | 535 | 709 |
| Phosphodiesterase | 16 | 0 (0.0%) | 0 (0.0%) | 449 | 888 |
| Protein-glutamine aminoacyltransferase | 7 | 0 (0.0%) | 0 (0.0%) | 693 | 817 |
| Cytochrome P450 | 103 | 0 (0.0%) | 0 (0.0%) | 503 | 561 |
| Aminoacyltransferase | 6 | 0 (0.0%) | 0 (0.0%) | 155 | 640 |

**Table S17. Sequence-length prevalence across enzyme families.**

Long proteins included several heavily studied targets, showing that the constraint is not restricted to rare proteins.

| ChEMBL target | Target name | Length (aa) | Activities | Family |
|---|---|---|---|---|
| CHEMBL203 | Epidermal growth factor receptor | 1210 | 28741 | kinase protein |
| CHEMBL2041 | Proto-oncogene tyrosine-protein kinase | 1114 | 26779 | kinase protein |
| CHEMBL2971 | Tyrosine-protein kinase JAK2 | 1132 | 24517 | kinase protein |
| CHEMBL1163125 | Bromodomain-containing protein 4 | 1362 | 23856 | regulator reader |

| | | | | |
|---|---|---|---|---|
| CHEMBL240 | Voltage-gated inwardly rectifying potassium channel | 1159 | 20137 | channel VGC |
| CHEMBL2835 | Tyrosine-protein kinase JAK1 | 1154 | 19794 | kinase protein |
| CHEMBL279 | Vascular endothelial growth factor receptor | 1356 | 18759 | kinase protein |
| CHEMBL3553 | Non-receptor tyrosine-protein kinase | 1187 | 17171 | kinase protein |
| CHEMBL3130 | Phosphatidylinositol 4,5-bisphosphate-related target | 1044 | 15630 | transferase |
| CHEMBL2148 | Tyrosine-protein kinase JAK3 | 1124 | 14830 | kinase protein |
| CHEMBL4005 | Phosphatidylinositol 4,5-bisphosphate-related target | 1068 | 12507 | transferase |
| CHEMBL4296 | Sodium channel protein type 9 subunit | 1988 | 10798 | channel VGC |

| | | | | |
|---|---|---|---|---|
| CHEMBL3717 | Hepatocyte growth factor receptor | 1390 | 10440 | kinase protein |
| CHEMBL1865 | Protein deacetylase HDAC6 | 1215 | 10282 | regulator eraser |
| CHEMBL3267 | Phosphatidylinositol 4,5-bisphosphate-related target | 1102 | 9986 | transferase |
| CHEMBL5936 | Toll-like receptor 7 | 1049 | 9507 | receptor toll-like |
| CHEMBL2973 | Rho-associated protein kinase 2 | 1388 | 8999 | kinase protein |
| CHEMBL1862 | Tyrosine-protein kinase ABL1 | 1130 | 8807 | kinase protein |
| CHEMBL1075104 | Leucine-rich repeat serine/threonine-protein kinase | 2527 | 8233 | kinase protein |
| CHEMBL4409 | cAMP/cGMP phosphodiesterase-related target | 1055 | 8138 | phosphodiesterase PDE10 |

**Table S18. Most-studied targets with sequences longer than 1,000 residues.**

Only 21.1% of long targets fell within 100 residues of the limit; most required more than minor truncation.

| Sequence-length bin | Targets | Share of targets >1,000 aa |
|---|---|---|

| | | |
|---|---|---|
| 1,001–1,100 (minor truncation) | 291 | 21.1% |
| 1,101–1,500 | 562 | 40.7% |
| 1,501–2,000 | 249 | 18.0% |
| 2,001–5,000 | 257 | 18.6% |
| >5,000 | 22 | 1.6% |

**Table S19. Distribution of sequence lengths above the 1,000-residue input limit.**

These data establish the prevalence of the sequence-length issue and justify explicit routing. They do not establish that domain extraction improves prediction accuracy relative to another sequence representation. Accordingly, the sequence strategy is reported as an auditable input-handling safeguard rather than as a validated performance enhancement.
